# Scanning ultrasound-mediated memory and functional improvements do not require amyloid-β reduction

**DOI:** 10.1101/2023.06.16.545394

**Authors:** Gerhard Leinenga, Xuan Vinh To, Liviu-Gabriel Bodea, Jumana Yousef, Gina Richter-Stretton, Tishila Palliyaguru, Antony Chicoteau, Laura Dagley, Fatima Nasrallah, Jürgen Götz

**Affiliations:** Clem Jones Centre for Aging Dementia Research, Queensland Brain Institute, The University of Queensland, Brisbane, QLD, Australia; Queensland Brain Institute, The University of Queensland, Brisbane, QLD, Australia; Centre for Advanced Imaging, The University of Queensland, Brisbane, QLD, Australia; Proteomics Facility, The Walter and Eliza Hall Institute of Medical Research, Melbourne, VIC, Australia; Department of Medical Biology, The University of Melbourne, Parkville, VIC, 3052, Australia

## Abstract

A prevalent view in treating age-dependent disorders including Alzheimer’s disease (AD) is that the underlying amyloid plaque pathology must be targeted for cognitive improvements. In contrast, we report here that repeated scanning ultrasound (SUS) treatment at 1 MHz frequency can ameliorate memory deficits in the APP23 mouse model of AD without reducing amyloid-β (Aβ) burden. Different from previous studies that had shown Aβ clearance as a consequence of blood-brain barrier (BBB) opening, here, the BBB was not opened as no microbubbles were used. Quantitative proteomics and functional magnetic resonance imaging revealed that ultrasound induced long-lasting functional changes that correlate with the improvement in memory. Intriguingly, the treatment was more effective at a higher frequency (1MHz) than at a frequency within the range currently explored in clinical trials in AD patients (286 kHz). Together, our data suggest frequency-dependent bio-effects of ultrasound and a dissociation of cognitive improvement and Aβ clearance, with important implications for the design of trials for AD therapies.

**Summary:** The therapeutic effect of ultrasound on memory in AD mice leads to altered protein expression and improved functional connectivity in the absence of amyloid-β removal. Of two frequencies explored, the higher ultrasound frequency (1 MHz) is more effective.

## Introduction

Alzheimer’s disease (AD) is the leading cause of disability in the aging population, accounting for up to 80% of all cases of dementia. The overarching histopathological feature is that two key molecules, amyloid-β (AΒ) and tau, form insoluble aggregates. These develop into microscopically visible brain lesions known as Aβ plaques and neurofibrillary tangles that contain tau (Goedert and Spillantini, 2006). Both Aβ and tau dysfunction are believed to initiate and drive the degenerative process in AD, leading to a progressive impairment of memory, reasoning, and social engagement (Jack, 2022). A routinely used system to model aspects of AD and other dementias is represented by transgenic mice (Padmanabhan and Götz, 2023). Aβ plaque formation, together with memory impairment, has been reproduced in several transgenic lines including APP23 mice. This strain expresses the human amyloid precursor protein (APP) together with the K670N/M671L pathogenic mutation present in familial cases of AD (Sturchler-Pierrat et al., 1997).

Given the challenges of developing effective pharmacological treatments for AD and other brain diseases, alternative strategies including low-intensity focused ultrasound (FUS) are currently being explored (Blackmore et al., 2023). At high intensity, ultrasound is being used as a focused, incisionless, FDA-approved surgical tool for ablating brain tissue and thereby treating diseases such as essential tremor and tremor-dominant Parkinson’s disease (Meng et al., 2021). At low intensity, ultrasound is being investigated as neuromodulatory and/or blood-brain barrier (BBB) opening tool in multiple animal species and human study participants (Blackmore et al., 2023; Meng et al., 2021).

To achieve its bio-effects, ultrasound induces mechanical effects, which can be categorized into acoustic radiation forces and cavitation. Thermal effects are largely negligible at low ultrasound intensities. When ultrasound is delivered focally and interacts with intravenously injected microbubbles (FUS^+MB^), cavitation is augmented, leading to BBB opening. This causes an uptake of blood-borne factors, some of which with therapeutic effects (Meng et al., 2021). FUS^+MB^ is used by us in a scanning mode, SUS^+MB^, to treat the entire brain. Several research groups have thereby been able (i) to reduce amyloid plaque load in AD mouse models (such as APP23 mice), without co-injecting exogenous drugs (Jordao et al., 2013; Leinenga and Götz, 2015; Shen et al., 2020), and (ii) to improve memory functions (Jordao et al., 2013; Leinenga and Götz, 2015; Shen et al., 2020). Following treatment, Aβ was found to be taken up and cleared by microglial cells. This robust Aβ clearance was assumed to be the reason for the improved memory functions.

Ultrasound has also been explored without delivering microbubbles (termed by us FUS^only^ or SUS^only^), a modality whereby cavitation effects are largely absent and the effects of ultrasound are neuromodulatory, utilizing mechanical forces. Such a SUS^only^ treatment, however, proved insufficient to clear Aβ (Leinenga et al., 2019). Given that low-intensity ultrasound is increasingly being explored as a novel treatment option in AD patients, either with microbubbles (Epelbaum et al., 2022; Lipsman et al., 2018; Meng et al., 2019) or without microbubbles (Beisteiner et al., 2020), we asked whether the SUS^only^ treatment paradigm could improve spatial memory in APP23 mice, and whether Aβ reductions are required as one may assume because Aβ is the culprit causing the cognitive impairments in the first place. Here, we explored two frequencies: We used 1 MHz (in the following, we term this SUS^only^ paradigm HighF), a frequency routinely used in mice for the purpose of BBB opening (Leinenga and Götz, 2015; Nisbet et al., 2017) and via intracranially implanted transducers in AD patients (Epelbaum et al., 2022) We also used 286 kHz (terming this SUS^only^ paradigm LowF), this being a frequency that is within the range (210-650 kHz) at which transcranial human FUS^+MB^ treatments are generally being performed (Lipsman et al., 2018; Meng et al., 2019), because at 1 MHz, attenuation and aberration of the human skull are too high to achieve safe and efficacious bio-effects.

Interestingly, we found that repeated HighF treatments of APP23 mice, unlike LowF treatments, improved spatial memory in the active place avoidance (APA) behavior-testing paradigm. Contrary to the prevalent view that Aβ pathology (in addition to tau) drives the degenerative process and that for cognitive functions to improve, Aβ needs to be reduced, we found that the memory improvements occurred in the absence of Aβ reductions. Reduced neuronal connectivity may be responsible for the symptoms of AD. Resting-state functional magnetic resonance imaging (rsfMRI) connectivity studies in AD patients have shown that there is a dysfunction in network-level organization of the brain, including the default mode, salience, and limbic networks (Palop et al., 2006; Zhou and Seeley, 2014). We therefore also investigated whether ultrasound treatment achieves a long-lasting modulation of these dysfunctional brain networks and therefore represents an alternative therapeutic approach to AD. Our results revealed that SUS^only^, particularly at the 1 MHz frequency, induced long-term structural and functional changes in neurons and their connections, as revealed by quantitative proteomics and analysis of correlations between rsfMRI and behavior testing. Together, our data suggest a dissociation of cognitive improvement and Aβ clearance, with important implications for AD treatment strategies. They further suggest that the current clinical trials may not use optimal ultrasound parameters, with our data providing new insights for ultrasound treatment optimization.

## Results

### High-frequency SUS improves spatial memory performance

To investigate the effects of low-intensity ultrasound in the absence of microbubbles (i.e., no opening of the BBB, SUS^only^) in an animal model of AD, we used two types of transducers, one operating at 1 MHz and a second at 286 kHz, both achieving a similar pressure inside the skull of 0.57 MPa (Fig. 1A). The study comprised four treatment arms: untreated wild-type (WT) mice, and APP23 mice that either received 8 weekly treatments at 1 MHz in a SUS^only^ scanning mode (HighF), or 286 kHz in a SUS^only^ scanning mode (LowF) and sham (i.e. with the mice being anesthetized and placed under the transducer but without receiving an ultrasound treatment) (Fig. 1B). Prior to the first treatment the mice underwent a 5-day spatial memory and learning assessment in the APA test, followed by an allocation to the treatment arms based on their performance. Following treatment, the mice were retested using the 5-day APA spatial memory testing paradigm, and then underwent an fMRI assessment, after which they were sacrificed and the brains dissected for analysis, including label-free quantitative proteomics (Fig. 1B).

**Figure 1.**
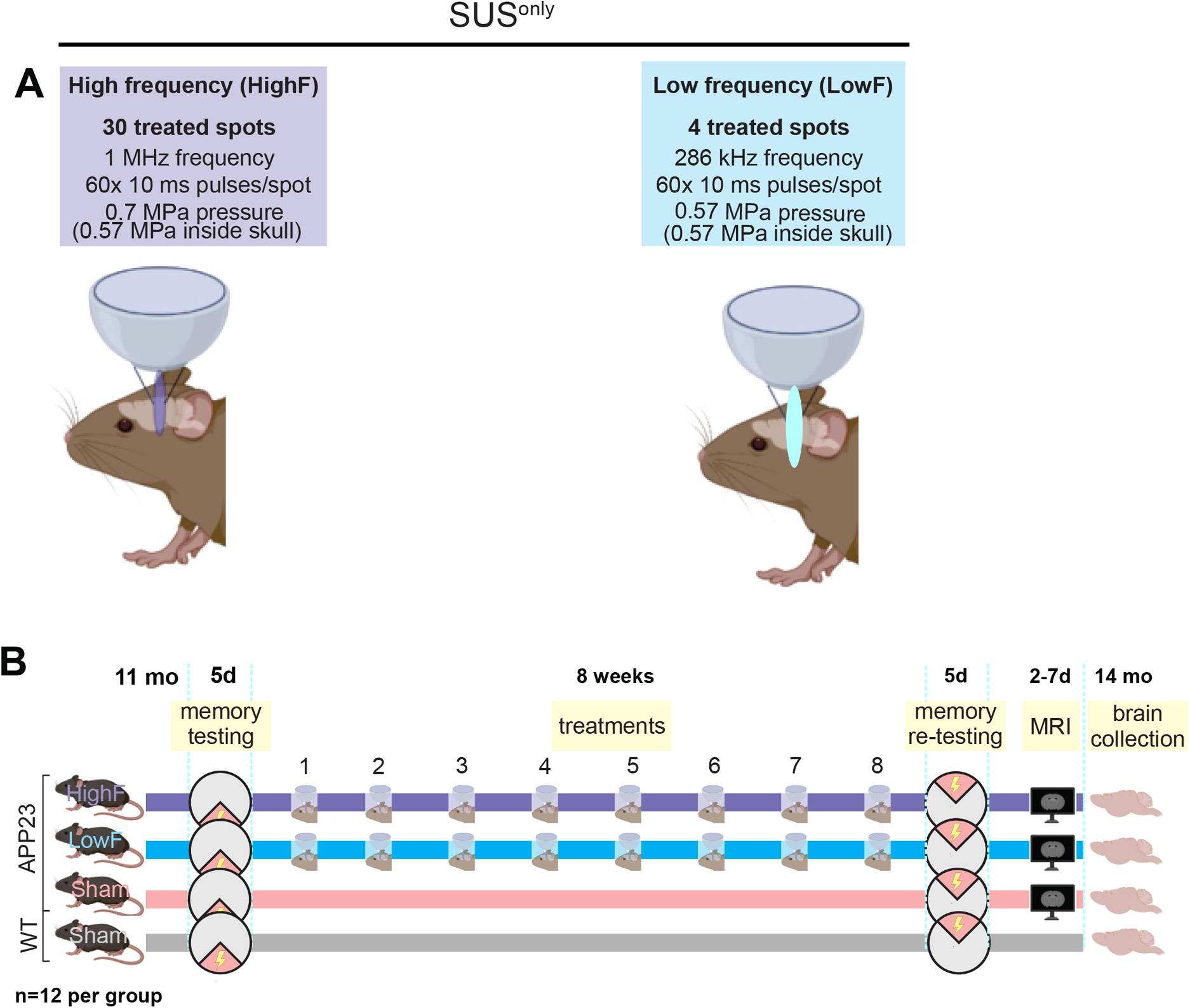
Study design. (A) APP23 mice aged 11 months were treated with scanning ultrasound (SUS^only^) at 1 MHz center frequency (HighF) or 286 kHz center frequency (LowF) with conditions arriving at the same pressure inside the skull accounting for the higher attenuation of high frequency ultrasound. Exposure of the whole brain was achieved by treating a 5×6 grid of spots with the HighF device, and 4 spots with the LowF device, taking into account the different −6 dB widths of the ultrasound focus at HighF (1 MHz, 1.5 mm) versus LowF (286 kHz, 6 mm). (B) APP23 mice and wild-type (WT) littermate controls were tested in the active place avoidance (APA) test over 5 days of training. APP23 mice then received eight once-per-week ultrasound treatments with either the HighF or LowF device. Controls received a sham treatment consisting of anesthesia and being placed under the ultrasound transducer, but the transducer was not turned on. WT mice were sham treated. Three days after the last ultrasound treatment, mice were retested in the APA retest in which the extra-maze cues were changed, the shock zone was placed in the opposite quadrant, and the arena rotated in a different direction. Between 2 and 7 days after the conclusion of the APA, mice received magnetic resonance imaging (MRI) scans and were sacrificed at the conclusion of the scans.

More specifically, we first examined 11 month-old APP23 mice and their wild-type littermates in the APA test, a hippocampus-dependent spatial learning paradigm in which the animals must use visual cues in order to learn to avoid a shock zone located in a rotating arena (Fig. 2A). This test is more suitable to probe memory functions than the Morris water maze, as aged mice are poor swimmers. The mice were first habituated to the arena in one 5-minute session the day before the first training day. The APA test then consisted of five training days with a single 10-minute training session each day.

**Figure 2.**
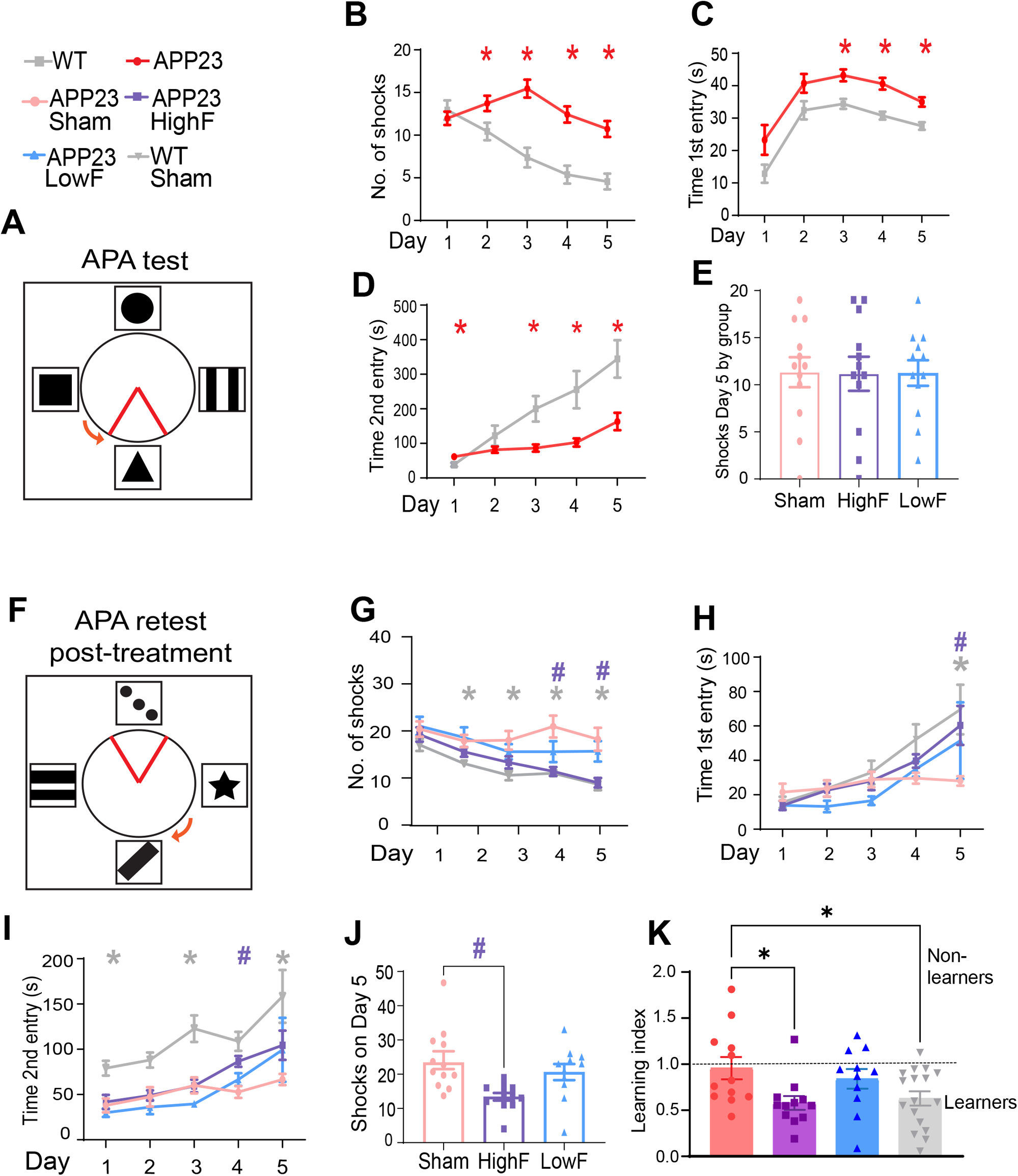
Improved spatial memory in APP23 mice in response to SUS^only^ at 1 MHz (HighF). (A) Schematic of the arena for the APA test, in which mice must use spatial cues to learn to avoid a shock zone within a rotating arena. APP23 mice show impaired performance compared to their WT littermates on the measures (B) number of shocks, (C) time to first entry of the shock zone, and (D) time to second entry to the shock zone. (Two-way ANOVA, * = p<0.05, APP23 compared to WT). (E) All mice were ranked based on their performance on the last day (day 5) of the APA and allocated to groups based on matching performance. (F) Following eight ultrasound treatments (1 MHz ultrasound (HighF), 286 kHz ultrasound (LowF) or sham), the mice were retested in the APA with the shock zone location, extra-maze cues and the direction of the arena rotation altered. There were significant differences between the treatment groups, such that compared to sham-treated APP23 mice, (G) HighF-ultrasound treated APP23 mice and sham-treated WT mice received fewer shocks, (H) had an increased time to first entry on day five (I), and had an increase in the time to second entry of the shock zone. (J) HighF-treated APP23 mice received significantly fewer shocks on day five of the APA retest compared to sham-treated APP23 mice. (Two-way ANOVA with follow-up Holm-Sidak test, # p<0.05 HighF compared to sham, * p<0.05 WT compared to sham-treated APP23 mice, # p<0.05 HighF-treated APP23 mice compared to sham-treated APP23 mice). (K) A learning index was calculated by dividing the number of shocks received on day five compared to day one of the APA retest, with better learning indicated by a lower ratio on the learning index. HighF mice demonstrated better learning on this index (One-way ANOVA with Holm-Sidak multiple comparisons test, * p<0.05).

In this first APA for group assignment, a two-way mixed repeated-measures ANOVA based on the number of shocks that were received revealed a significant effect of day of testing, indicating that learning had occurred ((*F*_4,188_) = 8.96, *p* < 0.0001). There was also a significant effect of genotype, with APP23 mice receiving more shocks than their wild-type littermates ((*F*_1, 47_) = 19.47, *p* < 0.0001) (Fig. 2B). Similarly, based on the measure of time to the first entry of the shock zone, there was a significant effect of day, with mice showing longer latencies to the first entrance as the number of training days increased ((*F*_4,188_) = 19.79, *p* < 0.0001). Wild-type mice exhibited longer latencies to enter the shock zone over the days of testing and there was a significant effect of genotype on time to first entry ((*F_1.47_*) *=* 13.13, *p* < 0.0001) (Fig. 2C). They also performed significantly worse on time to second entry as a measure of short-term memory, ((*F*_1,47_) = 24.16, *p* < 0.0001) (Fig. 2D). As the APA performance of the APP23 mice varied significantly, they were therefore assigned to each of the four treatment groups based on matching performance in terms of the number of shocks received on day 5 of the APA in order to reduce differences in performance between treatment groups (Fig. 2E).

Before retesting in the APA, mice were treated once-a-week for 8 weeks with either 1 MHz ultrasound (HighF), 286 kHz frequency ultrasound (LowF) or sham (sham) treatment. For the retest, the shock zone was shifted by 180°, the cues in the room were changed, and the arena was rotated in the opposite direction (Fig. 2F). To perform well in the retest, the mice needed to update their spatial learning in order to learn the new shock zone location, which requires significant cognitive flexibility. A two-way ANOVA with group as a between-subjects factor and day as a repeated measures factor revealed a significant effect of ultrasound treatment group on number of shocks received ((*F*_3,48_) = 11.27, p < 0.0001) (Fig. 2G). Follow-up multiple comparisons tests showed that HighF mice received significantly fewer shocks than sham mice on days 4 (p =0.004) and 5 (p = 0.008). In contrast to HighF-treated mice, LowF mice did not show a significant improvement in spatial learning ability compared to sham-treated mice. There was no significant effect of ultrasound treatment on time to first entry of the shock zone, which increased over day of training for all groups and revealed a trend towards an effect of both ultrasound treatments ((*F*_3,48_) = 2.36, p = 0.08) on time to first entry into the shock zone, with a follow-up multiple comparisons test showing that ultrasound-treated mice had a longer latency to enter the shock zone on day 5 of the APA task (p = 0.035) (Fig. 2H). There was a significant effect of group on the time to the second entry to the shock zone ((*F*_3,48_) = 12.03, p<0.0001) and here the HighF group showed a benefit on day 4 of the test (Holm-Sidak multiple comparisons test; p=0.01) (Fig. 2I). The number of shocks received on day 5 of the APA revealed that HighF mice received 43% fewer shocks than sham-treated APP23 mice (Holm-Sidak multiple comparisons test; p = 0.004) (Fig. 2J). We computed a learning index from the ratio of the number of shocks received on day one and day five of the APA retest, with a lower score indicating better learning. This learning index revealed significantly better learning in HighF treated mice compared to sham treated mice (One-way ANOVA followed by Holm-Sidak multiple comparisons test, p=0.027) (Fig. 2K). Together, these results demonstrate that APP23 mice exhibited an improvement in spatial memory when treated with HighF ultrasound, with LowF ultrasound only resulting in slight improvements.

### Improved memory functions occur in the absence of reductions in amyloid plaques and Aβ levels

We next examined whether the ultrasound treatments might reduce Aβ levels in brain by first performing Campbell-Switzer silver staining of sham, HighF and LowF treated APP23 mice for plaques (Fig. 3A-C). The analysis revealed no reduction in the total plaque area when the ultrasound treatment groups were compared to the sham-treated controls. We calculated the percentage area occupied by plaque for the cortex, in 5-10 sections per mouse, assessing plaque burden in a one-in-eight series of sections along the rostral-caudal axis starting from the anterior commissure and ending at the ventral hippocampus. There were no statistically significant differences when comparing sham and ultrasound treatment (one-way ANOVA ((*F*_2,36_)=1.87, p = 0.17). A t-test to compare HighF to sham revealed a slight increase in plaque burden, but this was not statistically significant (p = 0.16) (Fig. 3D). When the number of plaques with an area larger than 20 µm^2^ was quantified and normalized to the area of the cortex, no differences in plaque numbers were revealed (*(F*_2,36_) = 0.97, p = 0.39) (Fig. 3E).

**Figure 3.**
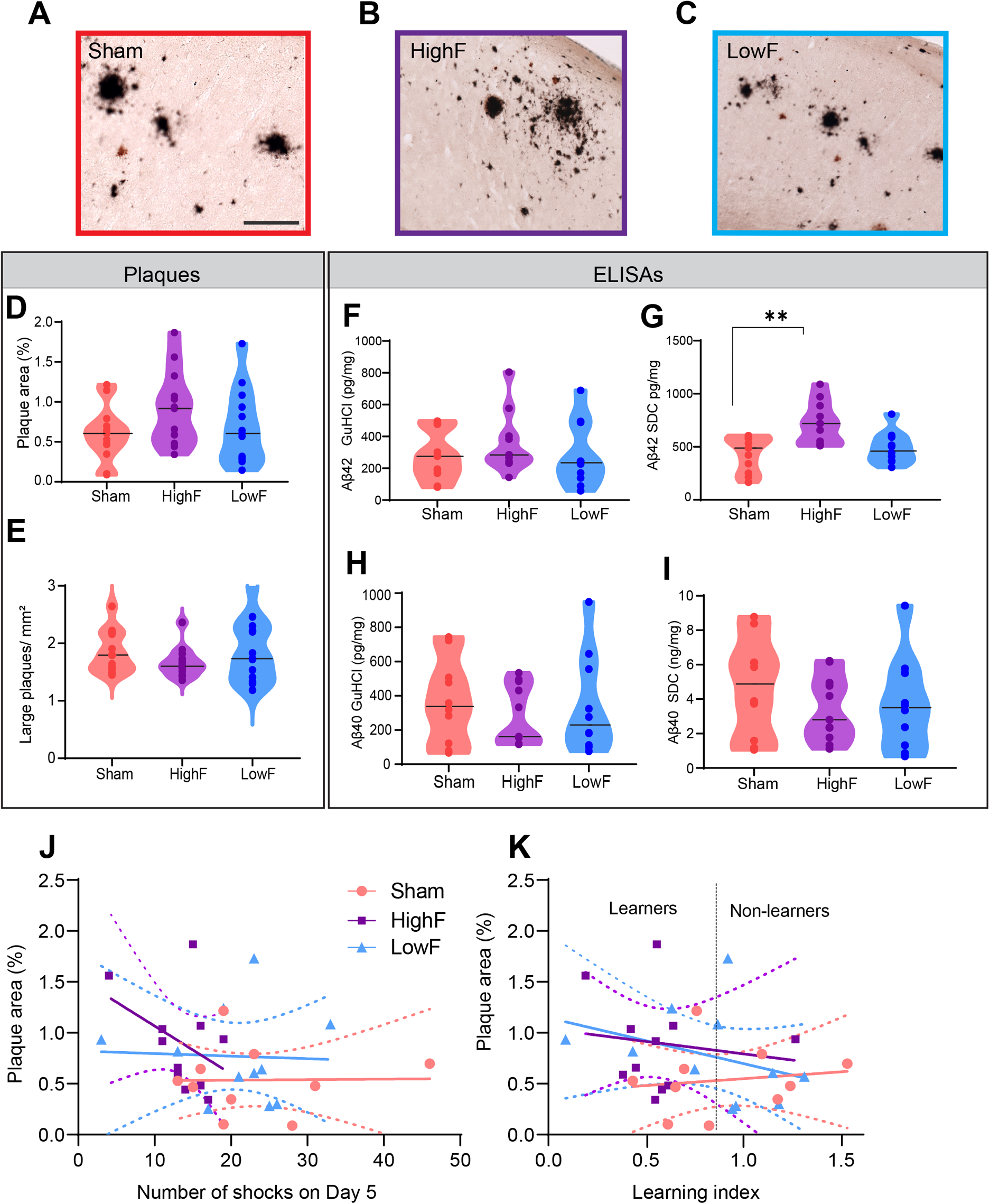
Repeated SUS^only^ treatments at either 1 MHz (HighF) or 286 kHz (LowF) does not reduce plaque burden and amyloid-β levels. (A) Plaque burden and morphology appeared similar when comparing sham-treated APP23 mice with (B) HighF-treated and (C) LowF-treated APP23 mice. The Campbell-Switzer silver staining method was used which stains both diffuse and compact plaques equally well. Plaque burden and number of large plaques were analyzed by automated thresholding in ImageJ. (Scale bar: 100µm). (D) Plaque burden expressed as % area was not significantly different between the three groups (one-way ANOVA p=0.16). (E) There was also no difference in the number of large plaques per mm^2^ of cortex. Cortical tissue was sequentially lysed to generate a detergent-soluble (SDC: sodium deoxycholate) fraction and a detergent-insoluble (GuHCl: guanidine hydrochloride fractions). (F) Enzyme-linked immunosorbency assays (ELISAs) for Aβ_40_ and Aβ_42_ revealed no difference in detergent insoluble Aβ_42_ levels, but (G) the detergent-soluble Aβ_42_ was higher in the HighF ultrasound treated group. (H) Levels of Aβ_40_ were not different between the groups in the detergent insoluble GuHCl fraction, or (I) in the detergent-soluble SDC fraction (I). Violin plots with the median value indicated with a line. (J) There was no correlation between amyloid-β plaque burden and the number of shocks the mice received on day 5 of the APA retest, and (K) no correlation between amyloid-β plaque burden and the learning index, calculated as the ratio of the number of shocks received on day one day five of the APA retest (Simple linear regression, slopes did not significantly differ from zero, dashed line is 95% confidence intervals).

As an additional assessment of amyloid pathology, we sequentially extracted the cortices of the mice to obtain a detergent-soluble fraction (SDC) containing soluble proteins and a detergent-insoluble fraction containing insoluble proteins (guanidine HCl) and measured the levels of the Aβ species, Aβ_40_ and Aβ_42_, by enzyme-linked immunosorbent assay (ELISA). Our results revealed no significant differences in the levels of Aβ_42_ in the insoluble fraction (Fig. 3F) (one-way ANOVA, p = 0.59) but a significant difference of Aβ_42_ in the soluble fraction (p = 0.001) (Fig. 3G). Levels of Aβ_40_ were not different between the treatment groups for the insoluble fraction (p = 0.71) (Fig. 3H) or the soluble fraction (Fig. 3I) (p = 0.51). We attempted to correlate plaque burden with performance in the APA for number of shocks on day five of the test (Fig. 3J), and the learning index (Fig. 3K), and found that there were no significant correlations. Together with the analysis of amyloid plaques, these findings reveal that improved memory function in the SUS^only^-treated APP23 mice occurred in the absence of amyloid reductions, and that spatial memory as measured in the APA test was not correlated with amyloid-β pathology including plaques.

### Ultrasound treatment alters the proteome of APP23 mice in a frequency-dependent manner

Given that ultrasound exerts pleiotropic effects on brain tissue, as reported recently in senescent WT mice (Blackmore et al., 2021), we performed label-free quantitative proteomic analysis on cortical lysates of APP23 mice subjected to either HighF, LowF or sham treatment. Principal component analysis (PCA) revealed that the proteomes of ultrasound-treated mice grouped together, away from the proteome of sham-treated mice, and that the HighF mice showed a relatively larger magnitude of changes in protein expression compared to the LowF animals, mirroring the differences seen in spatial memory functions (Fig. 4A).

**Figure 4.**
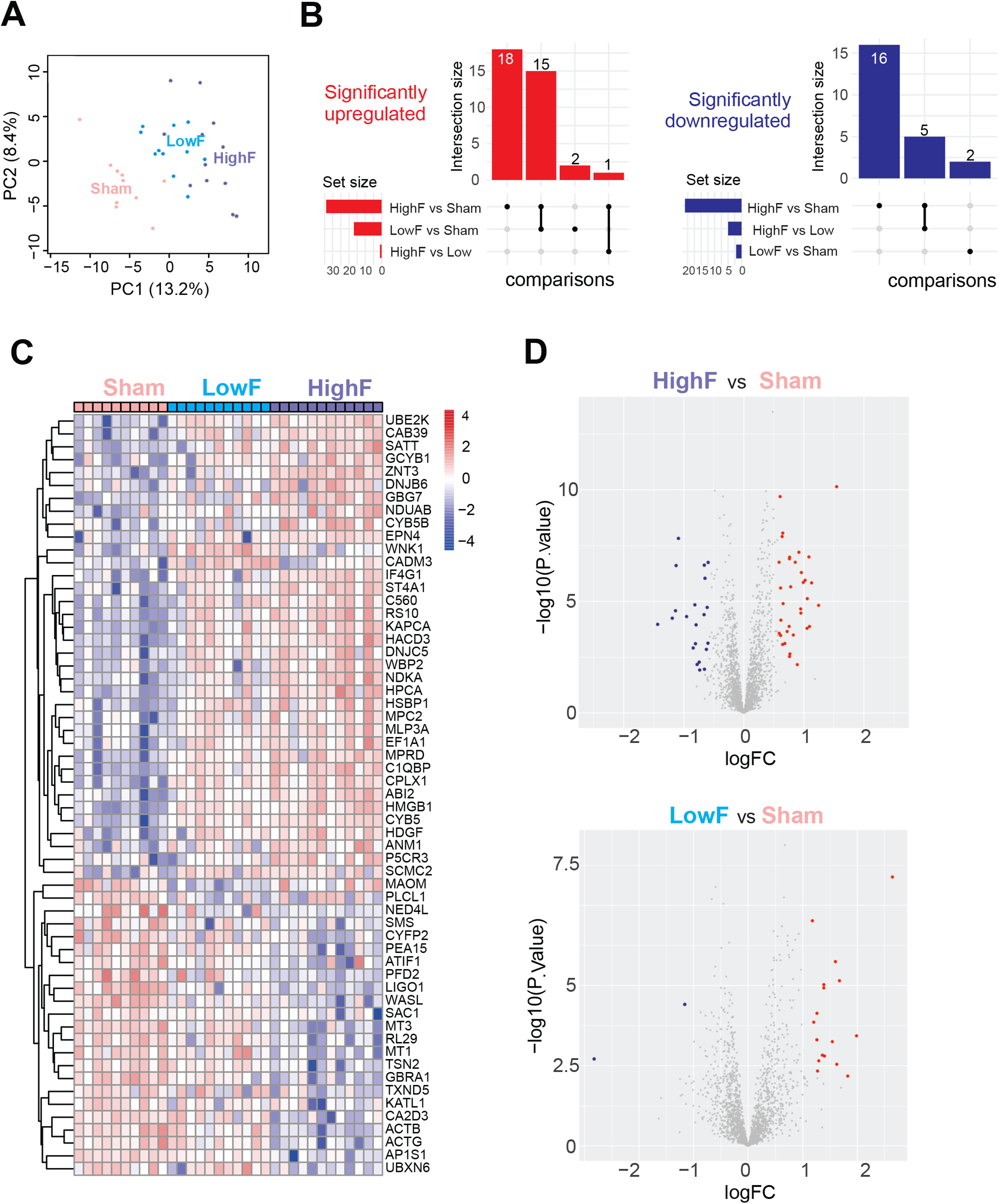
Proteomic analysis of the cortex of APP23 mice subjected to either HighF, LowF or sham treatment reveals treatment-dependent changes. (A) Principal component analysis (PCA) reveals the presence of a sham cluster independent of ultrasound treated samples, that show higher similarity, with both HighF and LowF ultrasound treatments clustering together. (B) Upset plot revealing the shared and unique number of proteins that were significantly upregulated (left) or downregulated (right) when compared between treatment regimes. (C) Heatmap of the top significant proteins (n=59) grouped by ultrasound treatment regime. (D) Volcano plot of differentially expressed proteins, displaying fold change (logFC, log_2_ scale) and *P* values (− log_10_ scale) between cuprizone HighF-treated vs sham-treated controls (top) and LowF-treated versus sham-treated controls (top).

Out of a total of 2,115 proteins identified by our proteomic pipeline, differential analysis revealed 15 upregulated proteins common to both the HighF and LowF mice compared to sham, and no proteins that were downregulated in both treatments when compared to sham (Fig. 4B). The same analysis identified 18 up- and 16 downregulated proteins that were specific to the HighF group compared with the sham-treated mice, and only 2 up- and 2 downregulated proteins in the LowF group compared with sham-treated mice (Fig 4B). Heatmap (Fig. 4C) and volcano plot representations of the differentially regulated proteins (Fig. 4D) revealed similar up- or down-regulation effects induced by the HighF versus sham treatment (32 significantly up- and 21 significantly downregulated proteins in volcano plots), whereas the LowF effect was, as expected, less impactful, but markedly skewed towards upregulation of protein expression (17 up- and 2 downregulated proteins versus sham) (Fig. 4D).

Our proteomic analysis led to the identification of four expression clusters based on their differentially expressed treatment-effect trajectory (Fig. 5 and Supplementary Fig. 1). Of these clusters, Cluster 1 (n=155 proteins) was characterized by increased expression in the LowF group and further enhanced by the HighF treatment (Fig. 5A). Cluster 1 was characterized by differential expression specific to Golgi vesicle transport, and establishment of protein localization to the membrane or to the cell periphery. Members of this cluster included SNX1 (member of the sorting nexin family), RAB5A (GTPase involved in endosome transport and found within different cellular components, including axon terminal boutons), and STX1b (involved in exocytosis of synaptic vesicles). Cluster 2 (n=147) (Fig. 5B) was represented by proteins that showed a marked ultrasound frequency-dependent downregulation of their expression. The functions associated with this cluster were represented by histone modification and response to hypoxia. Proteins which were downregulated in this cluster are represented by ACTB, SFPQ, NOS1 and MECP2. Interestingly, MECP2 is an intensely studied regulator of synaptic plasticity. Cluster 3 (n=186) was composed of proteins with upregulated expression following LowF treatment, but a more moderate expression increase induced by the HighF treatment (Supplementary Fig. 1A). This cluster was enriched in proteins associated with exocytosis, and axonal genesis and development. Interestingly, proteins belonging to this cluster included MAP1A and MAP6 (microtubule-associated proteins that contribute to axonal development and stabilization). Other members of the cluster were STX1A, SNX4 and CALM3, having functions related to exocytosis. Cluster 4 (n=173) was represented by proteins that showed a steep downregulation starting with the LowF treatment, but a moderate decrease after HighF treatment of mice (Supplementary Fig. 1B). The cluster was also found to be associated with axonogenesis and axonal development, as well as with vesicle organization. Proteins which were dysregulated in this cluster were associated with the action potential (GPD1L, GNA11, GNAQ) and localization of vesicles (RAB7A, DCTN2, MAP2).

**Figure 5.**
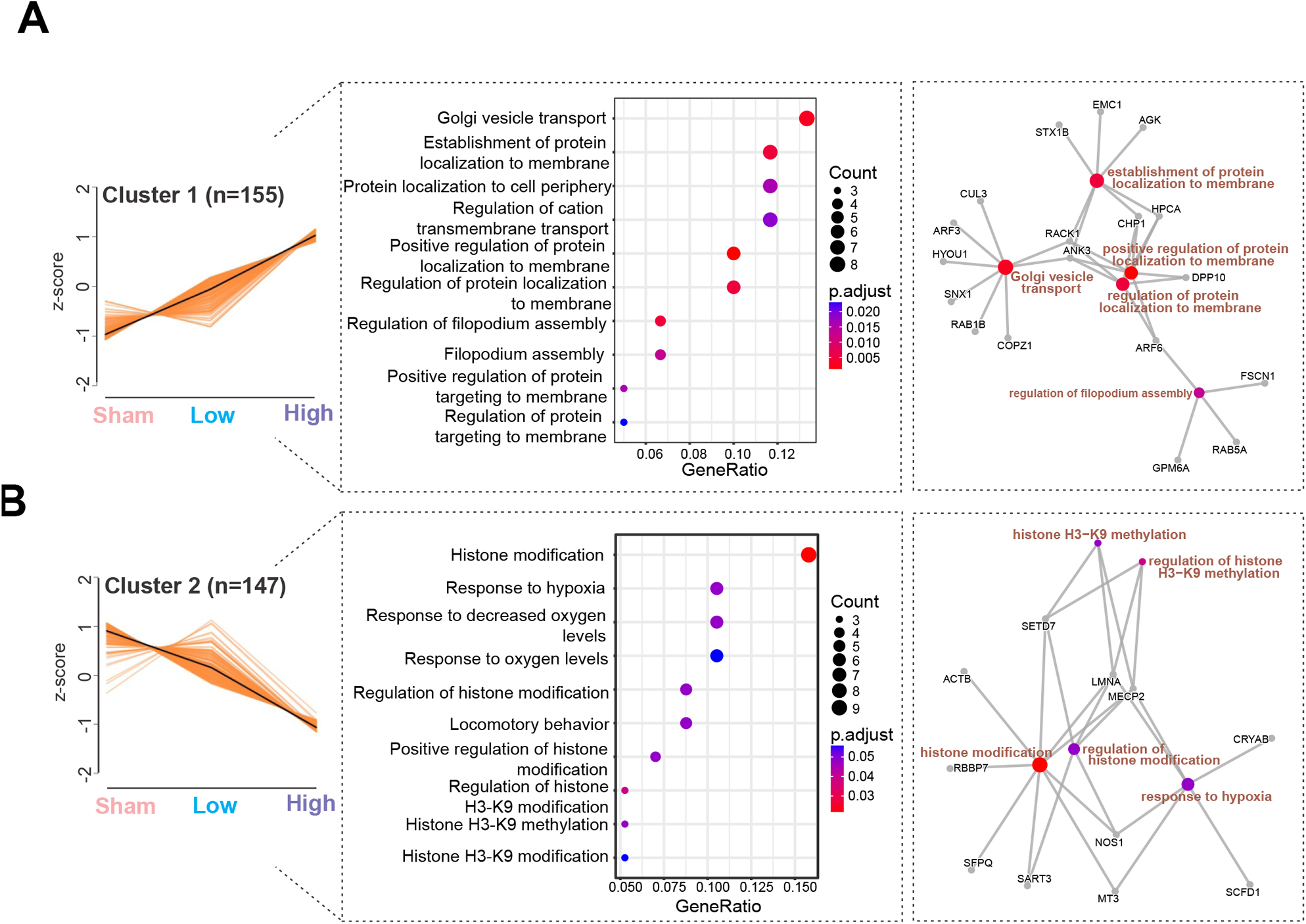
Bioinformatic analysis of proteomic data set identifies treatment-dependent differentially expressed clusters. Analysis of the proteomic data identifies expression patterns (left), top significant biological processes (middle) and expression networks (right) induced by the HighF and LowF ultrasound treatment. (A) Cluster 1 shows an ultrasound treatment-dependent increase in expression of proteins related to the Golgi vesicle transport and dynamics in membrane-associated proteins. (B) Cluster 2 is defined by an ultrasound treatment-dependent decrease in expression. The cluster is characterized by biological processes associated with histone modification.

Together, these findings not only indicate that HighF mice show a relatively larger magnitude of changes to the proteome compared to LowF and sham-treated animals, as expected, but they also provide a molecular explanation for the improvements seen in the spatial memory paradigm, through decreased transcriptional silencing of neuronal plasticity-related genes and an upregulation of the Golgi-related and synaptic vesicle secretory pathways.

### Ultrasound improves brain network connectivity as measured with rsfMRI

AD is associated with a widespread loss of both intra- and inter-network correlations as revealed by resting-state functional MRI (rsfMRI) (Brier et al., 2012), which has been reproduced in amyloid-depositing transgenic strains (Grandjean et al., 2016). Indeed, upon ultrasound treatment of amyloid-depositing APP23 mice, we generated functional connectivity matrices and found increased rsfMRI connectivity specifically in the hippocampal and salience networks in the HighF group compared to both sham and LowF mice (Fig. 6A-F). These findings indicate that the HighF treatment specifically improves connectivity in memory-related networks compared to other treatments and controls.

**Figure 6.**
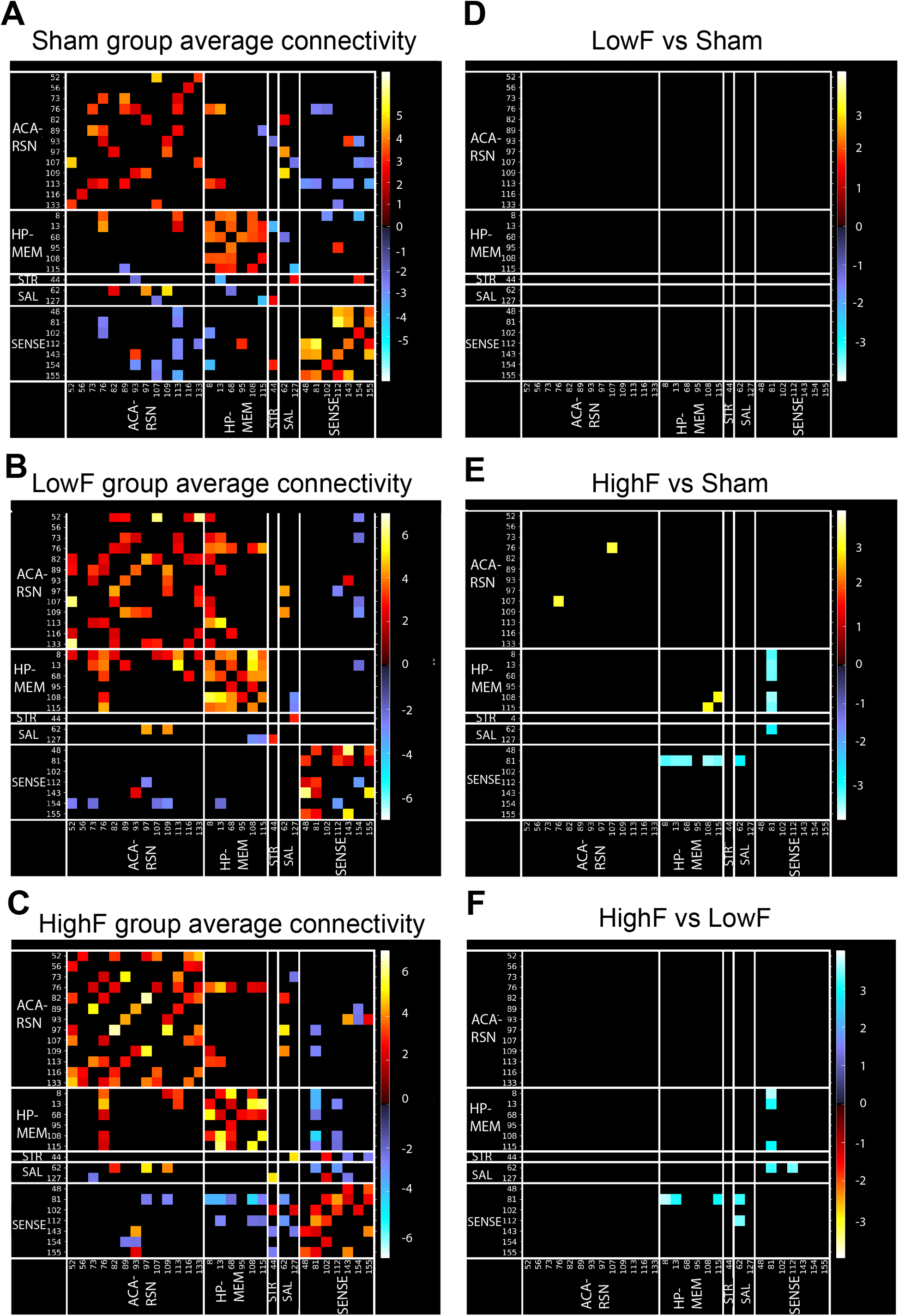
HighF ultrasound-treated mice show differences in averaged resting-state functional connectivity. Average resting state functional connectivity for selected brain regions for (A) sham-treated APP23 mice, (B) LowF-treated APP23 mice, (C) HighF-treated APP23 mice. No significant differences in functional connectivity were found when comparing the LowF- and sham-treated APP23 mice (D); however, HighF-treated APP23 mice showed significant differences in functional connectivity when compared to sham-treated APP23 mice (E), and when compared to LowF-treated APP23 mice (F). Color scaled by Z test statistics; non-black cells were defined as component-component connectivity deemed statistically significant. One sample t-tests, corrected for multiple comparison correction with False Discovery Rate, thresholded at Q < 0.05. Red–Yellow cells: positive correlation. Blue–Green cells: negative correlation.

Interestingly, stronger negative correlations were observed between the somatosensory network and the striatal and hippocampal networks in the HighF versus LowF and HighF versus sham groups (Fig. 6A-F). Negative correlations in resting state networks are an underexplored area yet evidence suggests that these opposing and anti-correlating regions can be interpreted as regions with opposing functions as the brain drifts between internal and external mental states (To et al., 2022). The enhanced negative correlations in the HighF group may indicate better switching ability across different mental states in these mice.

In contrast to the rsfMRI data, stimulus-evoked fMRI found no differences in ultrasound-treated compared to sham-treated APP23 mice (data not shown).

### Effect of ultrasound on brain microstructure

We next performed structural MRI and diffusion MRI imaging to investigate changes in brain volume and microstructure (Fig. 7A). Compared to the sham group, both the LowF and HighF groups had decreased local tissue volumes as measured by tensor-based morphometry (TBM). We then fitted the diffusion MRI data to the neurite-orientation density and dispersion index (NODDI) and found that the HighF treatment group showed a decreased orientation dispersion index (ODI) and a decreased neurite density index (NDI) mainly in the cortex (Fig. 7B). In contrast, the LowF group had increased ODI in the white matter tracts (Fig. 7C). Together, the alterations in the MRI diffusion metrics in the context of treatment in APP-related pathology may be associated with the observed enhanced neurocognition determined by APA spatial memory (Supplementary Fig. 2A,B). The decreased local tissue volume in the treated groups was also associated with increased isotropic diffusion fraction (fISO) predominantly in the neocortical areas and the anterior hippocampus, as well as increased volumes of the ventricles (Fig. 7B,C).

**Figure 7.**
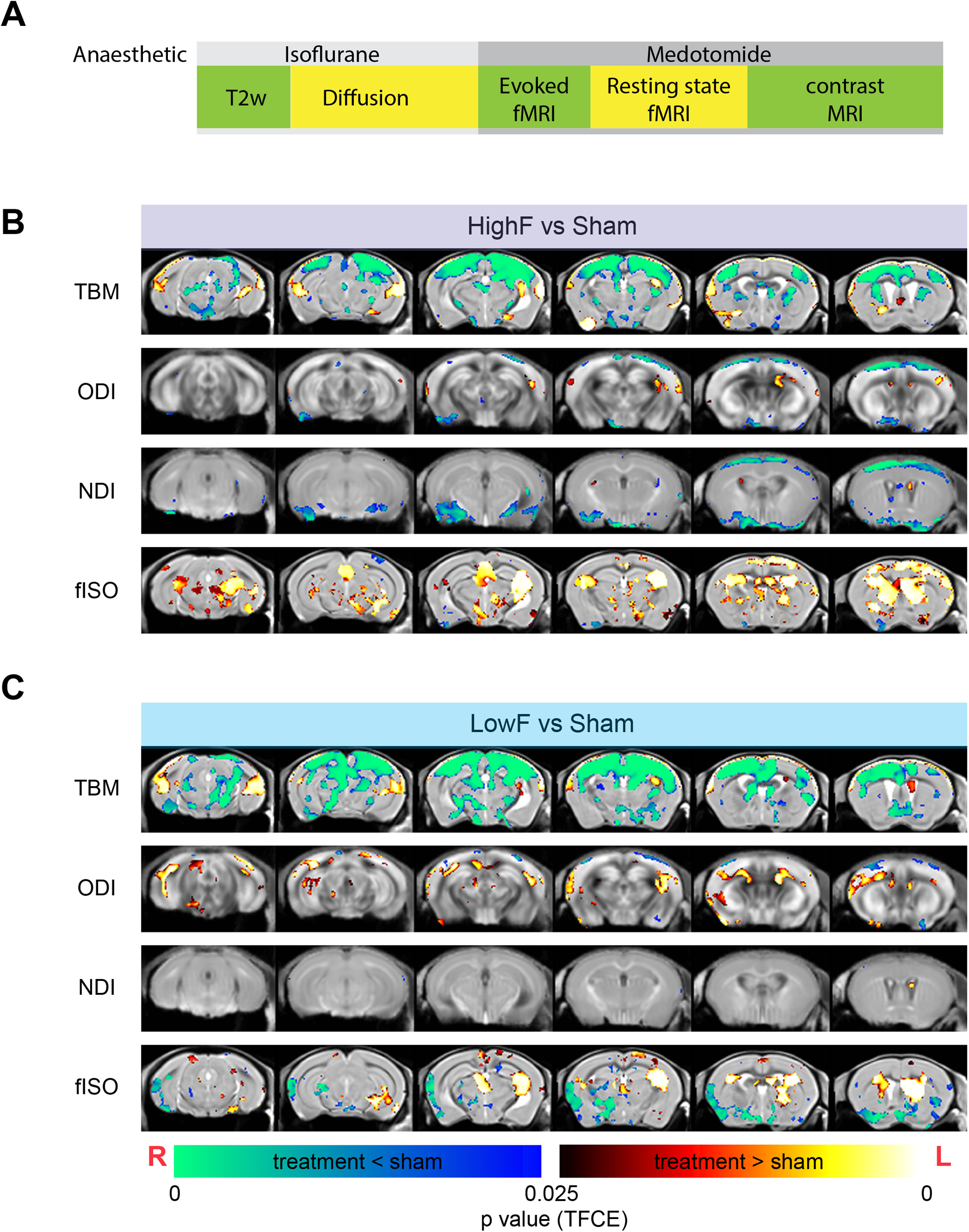
Changes to brain morphometry and NODDI diffusion measures by group. (A) Time-course of MRI and anesthesia. (B,C) Diffusion MRI reveals structural changes to the brain following HighF (B) and LowF ultrasound treatment (C). Red-yellow voxels show significant increase comparing treatment to sham, and blue-green voxels indicate significant reduction when comparing ultrasound treatment to sham. Tensor-based morphometry (TBM) visualized voxels that underwent statistically significant changes indicative of volumetric effects in HighF-treated APP23 mice. Neurite orientation dispersion and density imaging (NODDI) revealed changes in neurite density index (NDI), orientation dispersion index (ODI), and isotropic diffusion volume fraction (fISO) obtained from the diffusion model fitting. Similar changes were found in the LowF ultrasound-treated group (2 samples t-test results, implemented as randomized test of General Linear Model; statistical maps were corrected for multiple comparisons with Threshold-free Cluster Enhancement (TFCE) at P value < 0.05 (two-tailed). ODI = Orientation Dispersion Index, NDI = Neurite Density Index, fISO = isotropic diffusion volume fraction. Red anatomical orientation indicators: L = Left, R = Right).

We also performed dynamic contrast-enhanced MRI to measure cerebral blood volume; however, fitting of the Dynamic Susceptibility Contrast (DSC) data resulted in a large fraction of the datasets having cerebral blood volume (CBV) maps with predominantly negative values, even with leakage correction, indicating significant levels of BBB leakage in these animals (data not shown). This supports the notion that the BBB is impaired under many neurodegenerative conditions (Sweeney et al., 2019). Although DSC is not optimized for the detection of BBB integrity, we classified the animals into those giving feasible CBV value maps after DSC model fitting as having an intact BBB and those without feasible CBV maps (negative values) as having a leaky BBB. 55% of the sham mice had a leaky BBB by DSC MRI. 78% of the LowF mice and 88% of the HighF mice had a leaky BBB. A Chi-square test showed a statistically non-significant difference in the ratio of animals with leaky and non-leaky BBB across the treatment groups (p=0.8). This suggests that BBB integrity was not adversely impacted by the SUS treatment, and that BBB integrity is impaired in the majority of APP23 mice.

#### Resting state functional connectivity changes correlates with behavioral improvements in ultrasound-treated APP23 mice

We next asked whether the increased functional connectivity observed in ultrasound-treated mice is closely linked to the improvements in memory performance. Therefore, we used a group-independent component analysis to identify a subcortical memory network (To and Nasrallah, 2021b) comprising the posterior hippocampus, thalamus and the retrosplenial-anterior cingulate cortex, and assessed the connectivity strength in this network correlated with APA performance (Fig. 8A). We found a significant correlation between connectivity of the network and learning ability in the APA test, in particular the communication between the anterior thalamus and the retrosplenial cortex was associated with better learning in ultrasound-treated APP23 mice (grouping HighF and LowF) (Fig. 8B). Connectivity of this memory circuit was significantly correlated with performance in the APA test as determined by the learning index (Fig. 8C). We also measured the strength of the default mode network connectivity and ran the analysis but found that this network’s connectivity did not correlate with spatial learning performance (data not shown).

**Figure 8.**
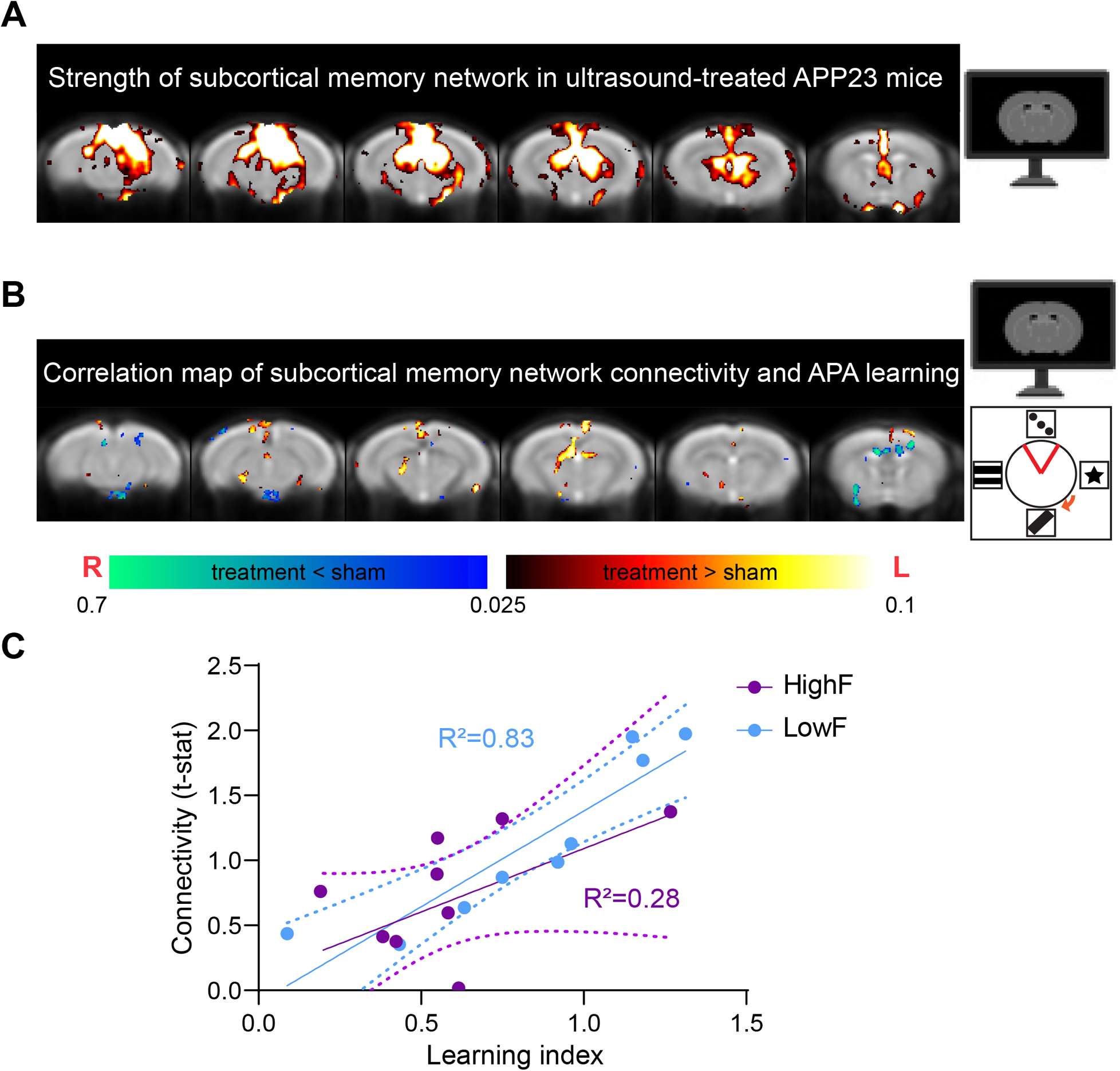
Functional connectivity is increased in HighF and LowF treated mice and correlates with memory performance in the APA test. (A) A subcortical memory network was identified using ICA analysis involving the posterior hippocampus, thalamus and the retrosplenial-anterior congulate cortex which is involved in spatial learning. (B) In ultrasound-treated mice, performance in the APA memory test was positively correlated with the subcortical memory network connectivity, particularly connectivity between the anterior thalamus and the retrosplenial cortex (Heat map indicates voxels with p <0.05 correlation). (C) Linear regression of functional connectivity in the subcortical memory circuit (t-statistic) and the learning index revealed a significant correlation in LowF treated mice (R^2^=0.83, p = 0.0006), and a correlation in HighF treated mice (R^2^=0.28, p = 0.1).

We further performed a correlation analysis of the diffusion metrics, NDI and ODI in the APA test and found that NDI was positively correlated with the number of shocks in the HighF group (Supplementary Fig. 2A). ODI was also positively correlated with the number of shocks (Supplementary Fig. 2B). Together, these data suggest that changes in the brain detected with fMRI or diffusion MRI correlate with spatial learning, possibly indicating that the detected alterations could reflect increased plasticity in response to ultrasound treatment which might lead to improvements in spatial memory functions.

#### Comparing (SUS^only^) HighF-mediated cognitive improvement in APP23 mice with those obtained previously using SUS^+MB^ that achieved BBB opening

We next compared the current data with those obtained in two previously published studies in which we had reported improved spatial memory in the APA paradigm using SUS^+MB^ (Leinenga and Götz, 2015; Leinenga et al., 2021). Our comparative post hoc analysis suggests that the effect of ultrasound on brain functions can be dissociated from amyloid reductions facilitated by BBB opening (Fig. 9A). Together, this reveals that SUS^only^, in particular the HighF paradigm, improved memory function, increased neuronal connectivity and altered the proteome in APP23 mice, without lowering Aβ levels (Fig. 9B).

**Figure 9.**
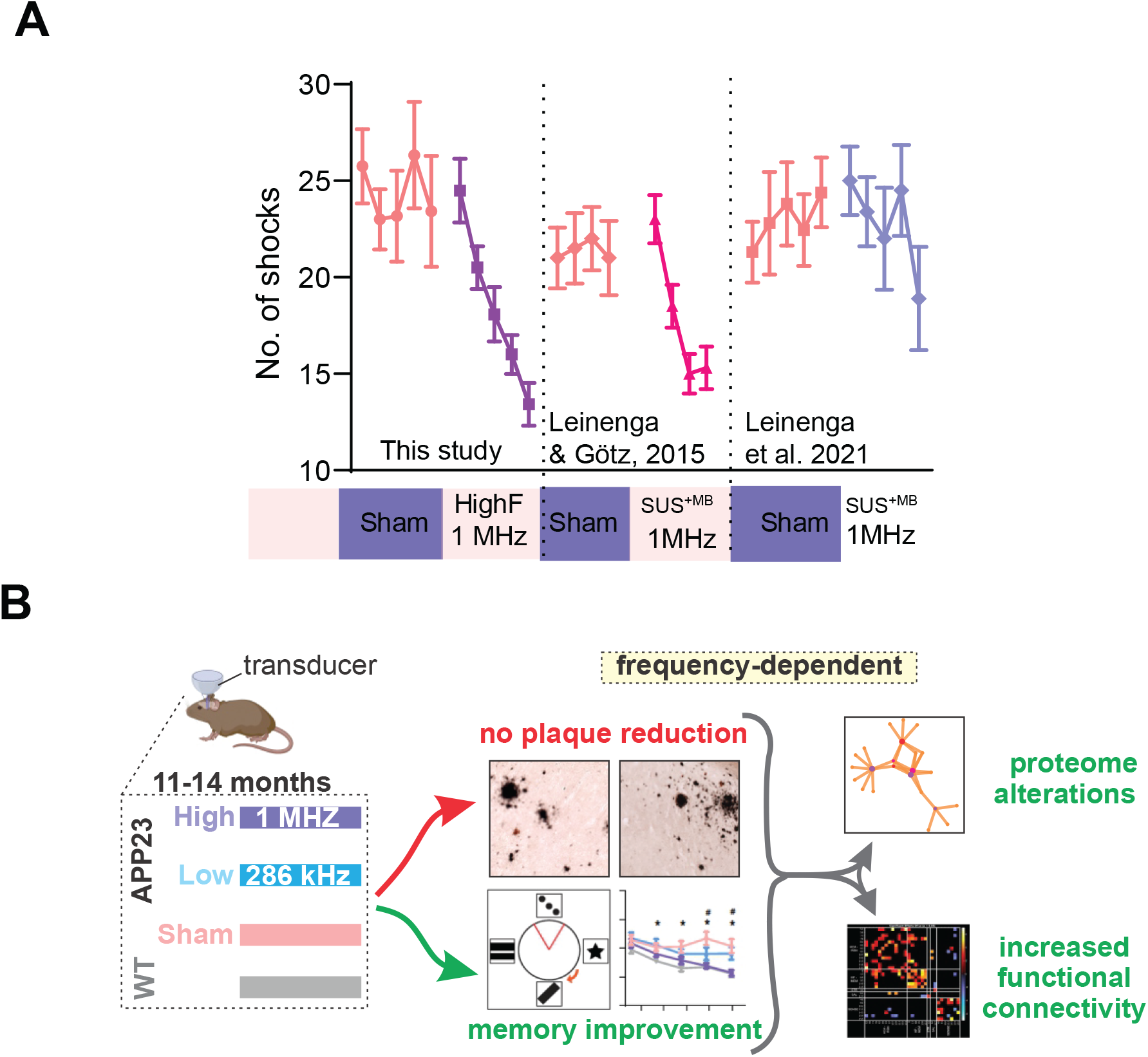
Effects of ultrasound (SUS only) treatment in APP23 mice. (A) The magnitude of the improvement in learning and memory after ultrasound treatment is equivalent to those obtained in APP23 mice treated with ultrasound plus microbubbles that opens the BBB (SUS^+MB^) reported in two earlier studies. (B) Improved memory, enhanced functional connectivity and altered proteome in APP23 mice in the absence of Aβ reduction after repeated treatments with scanning ultrasound without microbubbles (SUS^only^) at 1 MHz (HighF) and to some extent, 286 kHz (LowF).

## Discussion

In AD, memory deficits are assumed to be a consequence of the accumulation of Aβ (and tau) aggregation (Polanco et al., 2018; Selkoe and Hardy, 2016). Previously, we had revealed in APP23 mice with an Aβ pathology that ultrasound application in combination with the intravenous injection of microbubbles (SUS^+MB^) both safely and transiently disrupted the BBB; this led to amyloid clearance through increased microglial internalization, as well as memory improvements (Leinenga et al., 2022; Leinenga and Götz, 2015; Leinenga et al., 2021). The assumption at the time was that memory or functional improvements are contingent on reducing Aβ.

Here, we show that therapeutic ultrasound applied without microbubbles (SUS^only^), but with parameters shown previously to improve cognition in aged wild-type mice (Blackmore et al., 2021), improved memory function, increased neuronal connectivity and induced long-lasting changes in the proteome in APP23 mice, without lowering Aβ levels (Fig. 9A). Functional connectivity changes in a subcortical memory network were increased in ultrasound-treated mice and were correlated with improved performance in the APA spatial learning test. Changes to the neuronal proteome and brain microstructure were also observed and they correlated with ultrasound frequency in the case of proteomics and with memory performance in the case of brain microstructure as measured by diffusion MRI. In contrast, memory performance was not correlated with amyloid pathology. Together these findings might indicate that the symptomatic impairments seen in APP23 mice (and in human AD) can be targeted without necessitating the removal of amyloid, but through increasing neuronal function and connectivity. The improvements in memory and functional connectivity may be due to changes in synaptic protein expression and alterations in tissue microstructure, blood flow, and free water diffusion resulting from the ultrasound treatment. It has been shown previously that levels of synaptic proteins can predict the rate of cognitive decline in human AD patients when measured post mortem (Bereczki et al., 2016) or with positron emission tomography (PET) (Mecca et al., 2020). Interestingly, the reduced brain volume detected by T2-weighted MRI in the ultrasound-treated mice is consistent with other studies that have shown significantly increased brain volume loss in AD patients in clinical trials (Barkhof and Knopman, 2023). It has been speculated that amyloid plaque removal may explain the volume loss, but this is unlikely the reason given the small volume of brain occupied by plaques compared to the magnitude of changes found on MRI. How changes to brain microstructure detected in our study relate to changes in cognition could be subject of further studies, but we note that it occurred with both LowF and HighF ultrasound stimulation, although the latter had a greater effect on memory.

The SUS^only^ neuromodulation employed here improves memory in AD mice and in senescent wild-type mice (Blackmore et al., 2021). This finding has some similarities to other brain stimulation methods (such as repetitive transcranial magnetic stimulation) which have been found to improve memory function in healthy humans, specifically by stimulating regions of the prefrontal cortex (Luber and Lisanby, 2014). There is also evidence that brain stimulation can improve cognitive abilities in patients with AD, and clinical trials are underway (Menardi et al., 2022). Ultrasound is likely to have both overlapping and distinct mechanisms of action to electrical and magnetic stimulation, as it exerts a mechanical force on brain tissue and mediates its effects through the activation of mechanosensitive ion channels on neurons and glia (Oh et al., 2020; Yoo et al., 2022).

To the best of our knowledge, our study for the first time reveals frequency-specific effects of ultrasound on cognition and connectivity in mice. 1 MHz (HighF) was more effective than 286 kHz (LowF) in our study, possibly because of the increased acoustic radiation force exerted at the HighF condition. Another study reported that higher-frequency ultrasound has larger neurostimulatory effects on neurons in the retina (Menz et al., 2019). It is assumed that mechanical stimulation of neurons has a role in the effect of ultrasound, given that several different mechanosensitive ion channels are expressed by neurons that can respond to ultrasound (Blackmore et al., 2023). Magnetic resonance acoustic radiation force imaging (MR-ARFI) has shown that because absorption of ultrasound increases with frequency, it results in larger displacements in brain tissue (Hertzberg et al., 2010; Phipps et al., 2019). This increased absorption of high frequency ultrasound may explain the larger bioeffects when APP23 mice are treated at 1 MHz as compared to 286 kHz.

Along with improved cognitive function, we also detected changes in functional connectivity as measured by MRI. Interestingly, reduced functional connectivity determined by fMRI has been reported for 5xFAD mice (Kesler et al., 2018) and in a rat model of AD (Munoz-Moreno et al., 2018). We revealed that in addition to cellular level alterations induced by ultrasound, brain circuits may also be affected as reflected by changes in functional connectivity. Others have shown long-lasting effects (for 45 minutes) in primates after stimulation with 500 kHz ultrasound (Munoz et al., 2022). FUS^only^ further alters neuronal circuits and behaviors in macaques for up to one hour after stimulation (Verhagen et al., 2019). The long-term changes in functional connectivity occurring in our study two weeks after the last ultrasound treatment may be a consequence of alterations in the neuronal proteome that may underlie the improvements in cognitive function.

We found that ultrasound stimulation caused changes to the neuronal proteome lasting up to two weeks post-treatment, with the dysregulated clusters being tightly related to synaptic vesicle function, exocytosis, axonogenesis and neuronal signaling. The proteome changes we identified were both frequency dependent and independent, pointing to the mechanisms by which the SUS^only^ treatment could exercise its effects on the AD brain. Indeed, the HighF treatment induced an elevation in Golgi vesicle transport and membrane dynamics-associated processes and a marked decrease in histone methylation-associated processes. Together with the behavioral and MRI data, our proteomics analysis thus provides mechanistic evidence that could explain the HighF treatment effects: profound changes in chromatin organization and removal of transcriptional silencing through a decrease in the H3K9 methylation, simultaneous with increased secretory activity, and altogether leading to increased network connectivity and improved cognitive functions.

A limitation of our study is given that APP23 mice are characterized by premature lethality during the first 4-6 months of age which presents a challenge to breeding planning (Ittner et al., 2010), that the experimental groups were comprised of 75% male and 25% female mice; however, the ratio of males to females was the same for all treatment groups. Our study could therefore not ascertain a possible difference between male and female mice in their response to ultrasound. A second potential limitation of our study is that the treatment protocol with weekly treatments for two months was designed to be similar to a clinically relevant protocol. We did not assess the effect of ultrasound in real-time during ultrasound neurostimulation as we were interested in the amelioration of clinical symptoms rather than studying the online effects of ultrasound, and the immediate effects of ultrasound application or other timepoints remains to be determined.

In an AD context, ultrasound has previously been used to achieve concomitant BBB opening (FUS^+MB^) in mouse models (Leinenga and Götz, 2015; Xhima et al., 2021), and in a growing number of clinical trials enrolling small numbers of AD patients (Lipsman et al., 2018). A clinical trial has also been conducted in AD patients without microbubbles and using ultrasound shockwaves (Beisteiner et al., 2020). As reviewed recently, it is still incompletely understood how to best utilize ultrasound to achieve therapeutic outcomes (Blackmore et al., 2023). The current study reveals in the context of AD that SUS^only^ can differentially induce long-term functional improvements in the brain, as shown by quantitative proteomics and functional magnetic resonance imaging. Importantly, these improvements occurred in the absence of Aβ reductions. Our study therefore implies that cognitive improvement and Aβ clearance can be dissociated, with important implications for treatment strategies. We conclude that our findings are relevant for the rapidly growing space of therapeutic ultrasound, whether used as a neuromodulatory or BBB-opening tool for drug delivery.

## Materials and Methods

### Study design

APP23 mice express human APP751 with the Swedish double mutation (KM670/671NL) under the control of the neuron-specific mThy1.2 promoter. As the mice age, they exhibit memory deficits and amyloid plaque formation (Ittner et al., 2010). In this study, 12 months-old APP23 mice were assigned to three treatment groups: sham (N = 12), 1 MHz ultrasound (N = 12, HighF), and 286 kHz ultrasound (N=11, LowF). Assignment to treatment groups was based on matching performance of spatial memory (number of shocks) on day 5 of the APA test. APP23 mice were ranked from those receiving the fewest shocks to those receiving the most shocks on day 5 and were assigned to the three APP23 ultrasound treatment groups in rank order. A group of wild-type mice (N = 12) was included as fourth group. Following eight weekly treatments, mice underwent an APA re-test over five days, followed by an MRI analysis and the collection of brains for a proteomic, biochemical and histological analysis (Fig.1).

### Animal ethics

All animal experimentation was approved by the Animal Ethics Committee of the University of Queensland (approval number QBI/554/17).

### Ultrasound equipment

For the 286 kHz ultrasound application, a H-117 transducer was used (Sonic Concepts Bothell WA USA). This transducer has a diameter of 64 mm, a radius of curvature of 63 mm, and an internal opening of 20 mm diameter. For the 1 MHz ultrasound application, an integrated focused ultrasound system was used (Therapy Imaging Probe System, TIPS, Philips Research). This system consists of an annular array transducer with a natural focus of 80 mm, a radius of curvature of 80 mm, a spherical shell of 80 mm with a central opening of 31 mm diameter, a 3D positioning system, and a programmable motorized system to move the ultrasound focus in the x and y planes to cover the entire brain area. A coupler mounted to the transducer was filled with degassed water and placed on the depilated head of the mouse with ultrasound gel for coupling, to ensure propagation of the ultrasound to the brain.

### Ultrasound application

Mice were anesthetized with ketamine (90 mg/kg) and xylazine (6 mg/kg), and the hair on the head was shaved and depilated. The parameters for ultrasound delivery for the LowF condition were: 286 kHz center frequency, 0.57 MPa peak rarefactional pressure in situ (as estimated by needle hydrophone measurements obtained in water and assuming negligible attenuation through the skull), 10 Hz pulse repetition frequency (PRF), 10 ms pulse length (PL), and a 6 s sonication time per spot. Ultrasound was generated by a H-117 model transducer (Sonic Concepts) that was 64 mm diameter with a 63.2 mm radius of curvature and a 22 mm inner opening. The −6 dB focal zone was 6 mm (lateral) x 39.5 mm (axial). A motorized positioning system moved the focus of the transducer array to four spots spaced 6 mm apart in relation to the skull such that the entire brain of the mouse was exposed to ultrasound. The HighF condition was: 1 MHz centre frequency, 0.7 MPa peak rarefactional pressure (as estimated by needle hydrophone measurements in water for device calibration), 10 Hz PRF, 10 ms PL, and 6 s sonication per spot (Fig. 1). Ultrasound was generated by an eight-element annular array (TIPS system, Philips Research) with a diameter of 80 mm with an inner hole of 32 mm, an 80 mm radius of curvature. The focal zone of the array (−6 dB) was an ellipse of approximately 1.5 mm × 1.5 mm × 12 mm. The TIPS motorized positioning system moved the focus of the transducer array in a grid with 1.5 mm between individual sites of sonication so that ultrasound was delivered sequentially to the entire brain. The pressure level inside the brain was adjusted to be the equivalent for the HighF and LowF ultrasound groups (0.57 MPa derated) accounting for the higher, estimated 18% attenuation of 1 MHz ultrasound by the mouse skull (Choi et al., 2007).

### Assessment of amyloid plaques

For the assessment of amyloid plaque load, an entire one-in-eight series of coronal brain sections taken from the start of the anterior commissure to the ventral hippocampus of one hemisphere at 40 μm thickness was stained using the Campbell-Switzer silver staining protocol that stains both compact plaques and more diffuse amyloid fibrils but we did not distinguish these types in our analysis (Leinenga and Götz, 2015). Stained sections were mounted onto microscope slides and imaged with a 10x objective on a Metafer bright-field VSlide scanner (MetaSystems) using Zeiss Axio Imager Z2. Analysis of amyloid plaque load was performed using FIJI on all stained sections using ImageJ. A region of interest (ROI) was drawn around the cortex, and automated thresholding for plaques was applied to the ROIs, with the fill holes and despeckle functions applied to allow all plaques to be counted. A size threshold of 50 µm^2^ was applied to determine the number of plaques per area. No size thresholding was applied for the calculation of total plaque burden.

### Enzyme-linked immunosorbent assay for Aβ

Frozen cortices were homogenized in 10 volumes of a solution containing 1% sodium deoxycholate in 0.1M phosphate buffered saline with complete protease inhibitors, and homogenized by passing through 19 and 27 gauge needles. The samples were then centrifuged at 21,000×g for 90 min at 4 °C. The supernatant was retained as the detergent-extracted soluble Aβ fraction. The remaining pellets were resuspended in 10 volumes of 5 M guanidine HCl, sonicated, and centrifuged at 21,000×g for 30 min at 4 °C. The resultant supernatant was retained as the guanidine-extracted insoluble Aβ fraction. The concentrations of Aβ_40_ and Aβ_42_ were determined in brain lysates using ELISA kits according to the manufacturer’s instructions (human Aβ_40_ and Aβ_42_ brain ELISA, Invitrogen).

### Active place avoidance test

The APA task is a test of hippocampus-dependent spatial learning. We used a repeated APA paradigm, where mice were tested in the APA one time and the performance of each mouse was used to assign that mouse to one of four treatment groups. This was done by ranking all the mice based on their performance and assigning them to the four groups in order so that the APA performance of each treatment group was the same. Following this, mice received either sham, HighF or LowF treatment. Three days after the last treatment, they were retested to assess whether there was an improvement in APA performance. For each APA test, APP23 mice and non-transgenic littermate controls were tested over 6 days in a rotating elevated arena (Bio-Signal group) that had a grid floor and a 32-cm-high clear plastic circular fence enclosing a total diameter of 77 cm. High-contrast visual cues were present on the walls of the testing room. The arena and floor were rotated at a speed of 0.75 rpm, with a mild shock (500 ms, 60 Hz, 0.5 mA) being delivered through the grid floor each time the animal entered a 60-degree shock zone, and then every 1,500 ms until it left the shock zone. The shock zone was maintained at a constant position in relation to the room. Recorded tracks were analyzed with Track Analysis software (Bio-Signal). A habituation session was performed 24 h before the first training session during which the animals were allowed to explore the rotating arena for 5 min without receiving any shocks. A total of five training sessions were held on consecutive days, one per day with a duration of 10 min. After day 5 of the first APA (test), APP23 mice were divided into four groups with mice matched so that the performance (number of shocks) of the four groups of mice on day 5 of the task was the same, for the retest. Following eight once-a-week sham or ultrasound treatments, the mice underwent the APA test again (reversal learning). The retest was held in the same room as the initial test. However, the shock zone was switched to the opposite side of the arena, the visual cues were replaced with different ones, and the platform was rotated clockwise rather than counterclockwise. The number of shocks, numbers of entries to the shock zone, time to first entry, time to second entry, and proportion of time spent in the opposite quadrant to the shock zone were compared over the days of testing.

### Magnetic resonance imaging

Mice were imaged on a 9.4T (Bruker Biospin) MRI scanner between 8 and 12 days after the final (8^th^) high, low or sham treatment. Deep anesthesia was induced with 3% isoflurane in a 60/40 mixture of air/O_2_ at 1 l/min; the isoflurane concentration was reduced to 1.5–2 % for the remaining animal preparation time. Anesthetized mice were placed on the MRI-compatible cradle in a head-first, prone position and the head was fixed using ear bars and a bite bar to avoid movement. An intraperitoneal (i.p.) catheter was inserted and fixed temporarily for the infusion of medetomidine (Domitor, Pfizer) (Nasrallah et al., 2014). A 30-gauge tail vein catheter was inserted into a lateral tail vein for infusion of contrast agent (Magnevist, 0.5 mmol/kg). Subdermal electrodes were inserted near the 2nd and 4th digits on the left forepaw for mild electrical stimulations, which were delivered by a current source (Isostim A320, World Precision Instrument). The animals were monitored for respiration rate, pattern, and stability and rectal temperature (target 37 ± 0.5 °C) using an MRI-compatible rodent physiological monitoring system (Model 1030, SA Instruments).

Following proper positioning of the mouse inside the scanner magnet, an i.p bolus dose of 0.05 mg/kg medetomidine was given and continuous sedation was maintained with continuous medetomidine i.p infusion at 0.1 mg/kg/h. The isoflurane concentration was lowered gradually and maintained at 0.5% throughout the remainder of the experiment. T2-weighted, diffusion, BOLD and contrast-enhanced scans were obtained. The total time under anesthesia was approximately 2.5 h. At the end of the scan, an i.p bolus dose of 1.25 mg/kg atipemazole (for medetomidine reversal) was given and the animals were kept on a heated pad and monitored for recovery.

The sequence parameter details for the T2-weighted (T2w) structural, stimulus-evoked and resting-state functional, and multi-shell diffusion scans have been described previously (To and Nasrallah, 2021b). Briefly, T2w structural MRI scans were taken with a Turbo Rapid Acquisition with Refocused Echoes (TurboRARE) sequence with TR/TE = 7200/39 ms and a resolution of 0.1 × 0.1 × 0.3 mm. Stimulus-evoked and resting-state functional MRI scans were acquired with a 2D gradient-echo echo-planar-imaging (GE-EPI) sequence with TR/TE = 1000/14 ms, flip angle = 70°, and spatial resolution of 0.3 × 0.3 × 0.6 mm.

A Dynamic Susceptibility Contrast (DSC) sequence was acquired using a 2D gradient-echo echo-planar-imaging (GE-EPI) sequence with the following parameters: matrix size = 64 × 64, FOV = 19.2 × 19.2mm, 17 slices of 0.7 mm thickness and 0.1 mm slice gaps; giving an output spatial resolution of 0.3 × 0.3 × 0.8mm, TR/TE = 550/14 ms, flip angle = 35°, bandwidth = 3125 Hz/pixel, fat suppression = ON, FOV saturation (covering the head tissue inferior to the mouse’s brain) = ON, and navigator pulses = ON. The DSC sequence was started and allowed to run for 60 s before the contrast agent bolus was delivered; the DSC sequence was scanned with 982 volumes.

### Magnetic resonance imaging analysis

The preprocessing and image registration of the structural, diffusion, and functional MRI images were done based on detailed procedures as previously described (To and Nasrallah, 2021a). Briefly, structural images were corrected for bias field inhomogeneity and then normalized to a common space by an iterative image registration-template creation procedure, and the Jacobian images were used for a tensor-based morphometry analysis of structural differences. Diffusion MRI data were distortion and motion corrected, and fitted for the NODDI model, and also spatially normalized using a similar procedure to structural image. Orientation dispersion index (ODI), neurite density index (NDI), and fraction of isotropic diffusion (fISO) were obtained from the multi-shell diffusion MRI data. Functional MRI data were distortion and motion corrected, high-pass filtered at 0.01 Hz and also spatially normalized before being analyzed through a data-driven group independent component analysis (group ICA) using independent vector analysis – Gaussian and Laplacian source vecos (IVA-GL) into 65 independent components. Independent components were selected and grouped into larger networks (To and Nasrallah, 2021b). DSC MRI data were processed and spatially normalized to a common space using a process similar to that used for functional MRI data. Dynamic Susceptibility Contrast (DSC) data were fitted using the Dynamic Susceptibility Contrast MRI toolbox (https://github.com/marcocastellaro/dsc-mri-toolbox) with semi-automatic aterial input function selection and blood-brain barrier leakage correction (Boxerman et al., 2006).

### SWATH mass spectrometry

To identify changes in protein abundance following 1 MHz or 286 kHz ultrasound treatments compared to sham, total cortical protein extracts from 10-12 mice per group were analyzed by SWATH-MS. Briefly, 45 µg of protein from each sample was diluted in triethylammonium bicarbonate buffer and subsequently reduced with DTT, followed by alkylation with iodoacetamide. Samples were then digested with 80 ng of trypsin overnight, after which they underwent a buffer exchange to be resuspended in 2% acetonitrile, 0.1% formic acid. In order to form a unique peptide library, 5 µg from each of the samples were first pooled together and then fractionated via high pH reversed-phase high-performance liquid chromatography to form 18 fractions that were then analyzed via data-dependent acquisition (IDA).

For IDA, the sample (10 μL) was injected onto a reverse-phase peptide trap for pre-concentration and desalted with loading buffer, at 5 μL/min for 3 min. The peptide trap was then switched into line with the analytical column. Peptides were eluted from the column using a linear solvent gradient from mobile phase A: mobile phase B (95:5) to mobile phase A: mobile phase B (65:35) at 300 nL/min over a 60 min period. After peptide elution, the column was cleaned with 95% buffer B for 6 min and then equilibrated with 95% buffer A for 10 min before next sample injection. The reverse phase nanoLC eluent was subject to positive ion nano-flow electrospray analysis in an information dependant acquisition mode (IDA). In the IDA mode a TOF-MS survey scan was acquired (*m/z* 350-1500, 0.25 s) with the 20 most intense multiply charged ions (2+ - 5+; exceeding 200 counts per s) in the survey scan sequentially subjected to MS/MS analysis. MS/MS spectra were accumulated for 100 ms in the mass range *m/z* 100–1800 with rolling collision energy.

Data independent acquisition (SWATH) was performed as follows: Sample (10 μL, ∼ 2 μg) was injected onto a reverse-phase peptide trap for pre-concentration and desalted with loading buffer, at 5 μL/min for 3 min. The peptide trap was then switched into line with the analytical column. Peptides were eluted from the column using a linear solvent gradient from mobile phase A: mobile phase B (95:5) to mobile phase A: mobile phase B (65:35) at 300 nL/min over a 60 min period. After peptide elution, the column was cleaned with 95% buffer B for 6 min and then equilibrated with 95% buffer A for 10 min before next sample injection. The reverse phase nanoLC eluent was subject to positive ion nano-flow electrospray analysis in a data independent acquisition (SWATH). In SWATH mode, first a TOFMS survey scan was acquired (*m/z* 350-1500, 50 ms) and then, the 100 predefined *m/z* ranges were sequentially subjected to MS/MS analysis. MS/MS spectra were accumulated for 30 ms in the mass range *m/z* 350-1500.

Database searches for IDA data were performed as follows: The data files generated by IDA-MS analysis were searched with ProteinPilot (v5.0) (Sciex) using the ParagonTM algorithm in thorough mode. *Mus musculus* ‘uniprot-mus_musculus containing 17,119 proteins on July 4th 2022, was used for searching the data. Carbamidomethylation of Cys residues was selected as a fixed modification. An Unused Score cut-off was set to 1.3 (95% confidence for identification), and global protein FDR of 1%.

For SWATH extraction and quantitation, the local ion library was constructed using the 2D-IDAs. The library contains 4,448 proteins. Ion library and SWATH data files were imported into PeakView (v2.2) (Sciex). Protein peak area information in SWATH data were extracted using PeakView (v2.2) with the following parameters: Top 6 most intense fragments of each peptide were extracted from the SWATH data sets (75 ppm mass tolerance, 10 min retention time window). Shared and modified peptides were excluded. After data processing, peptides (max 100 peptides per protein) with confidence ≥99% and FDR ≤1% (based on chromatographic feature after fragment extraction) were used for quantitation.

### Bioinformatics analysis of SWATH-MS data

Data processing and analysis were performed using R (version 4.2.1). Proteins that were identified by two or more peptides were used in the analysis pipeline. A total of **2,115** proteins were included in the analysis. Protein intensities were log_2_-transformed. Normalization based on groups was performed using RUVIIIC (v. 1.0.19). Invariant proteins in all conditions (p > 0.5), with a coefficient of variation (CV%) < 2%, were chosen as negative controls for RUVIIIC normalization. Differential analysis was performed using limma (v. 3.52.4). A protein was determined to be significantly differentially expressed if the fold change was ≥ 1.5 or ≤ 0.5 and the false discovery rate (FDR) was ≤ 5% after Benjamini–Hochberg (H-B) correction.

Clustering analysis was performed based on mixed-effects models with Gaussian variables and a non-parametric cubic splines estimation approach implemented in TMixClust R package (v.1.18.0) GO (gene ontology) analysis and KEGG (Kyoto Encyclopedia of Genes and Genomes) pathways analysis were performed using clusterProfiler R-package (v. 4.4.4).

### Statistical analysis

Statistical analyses were conducted with Prism 9 software (GraphPad). Values were always reported as mean ± SEM. One-way ANOVA followed by the Holm-Sidak multiple comparisons test, or *t* test was used for all comparisons except APA analyses where two-way ANOVA with day as a repeated measures factor and group as a between subjects factor was performed, followed by the Holm-Sidak multiple comparisons test for simple effects to compare group performances on different days. The model assumption of equal variances was tested by Brown-Forsyth or Bartlett tests, and the assumption of normality was tested by Kolmogorov-Smirnov tests and by inspecting residuals with QQ plots. All observations were independent, with allocation to groups based on APA where mice were ranked on performance and assigned to one of the three groups (HighF, LowF, sham) in order of number of shocks on day 5 ranked from most to least shocks.

Spatially normalized MRI outputs, including Jacobian Index (obtained from T2w structural MRI image registration procedure), NODDI metrics (ODI, NDI, and fISO) and group ICA independent components’ spatial maps were analyzed as 2 sample t-tests using permutation inference for the General Linear model as implemented in FSL’s randomize (Winkler et al., 2014). The number of permutations was set to 10000, as recommended by a prior study (Dickie et al., 2015). The resulting statistical maps were corrected for multiple comparisons with mass-based FSL’s Threshold-free Cluster enhancement (TFCE) (Smith and Nichols, 2009) and thresholded at p < 0.05 (two-tailed). One and two-samples statistical tests were conducted on individual level independent components’ time courses using GIFT’s Mancovan toolbox for IC-IC functional network connectivity (FNC) and corrected for multiple comparisons with False Discovery Rate at q-value < 0.05 for one-sample t-tests and q-value < 0.1 for two-sample t-tests.

## Acknowledgements

We thank members of our team (Drs Rachel de las Heras and Jae Song) as well as Rowan Tweedale for critical reading of our manuscript.

## Author contributions

The study was designed by GL and JG. The experiments were performed by GL, XVT, GR-S and TS. The data were analyzed by GL, XVT, LGB, JY, AC and LD. The paper was written by GL, LGB, FN and JG. Funding was provided by JG.

## Data and materials availability

The data will be made available upon reasonable request.

## Supplementary figures

**Supplementary Figure 1.**
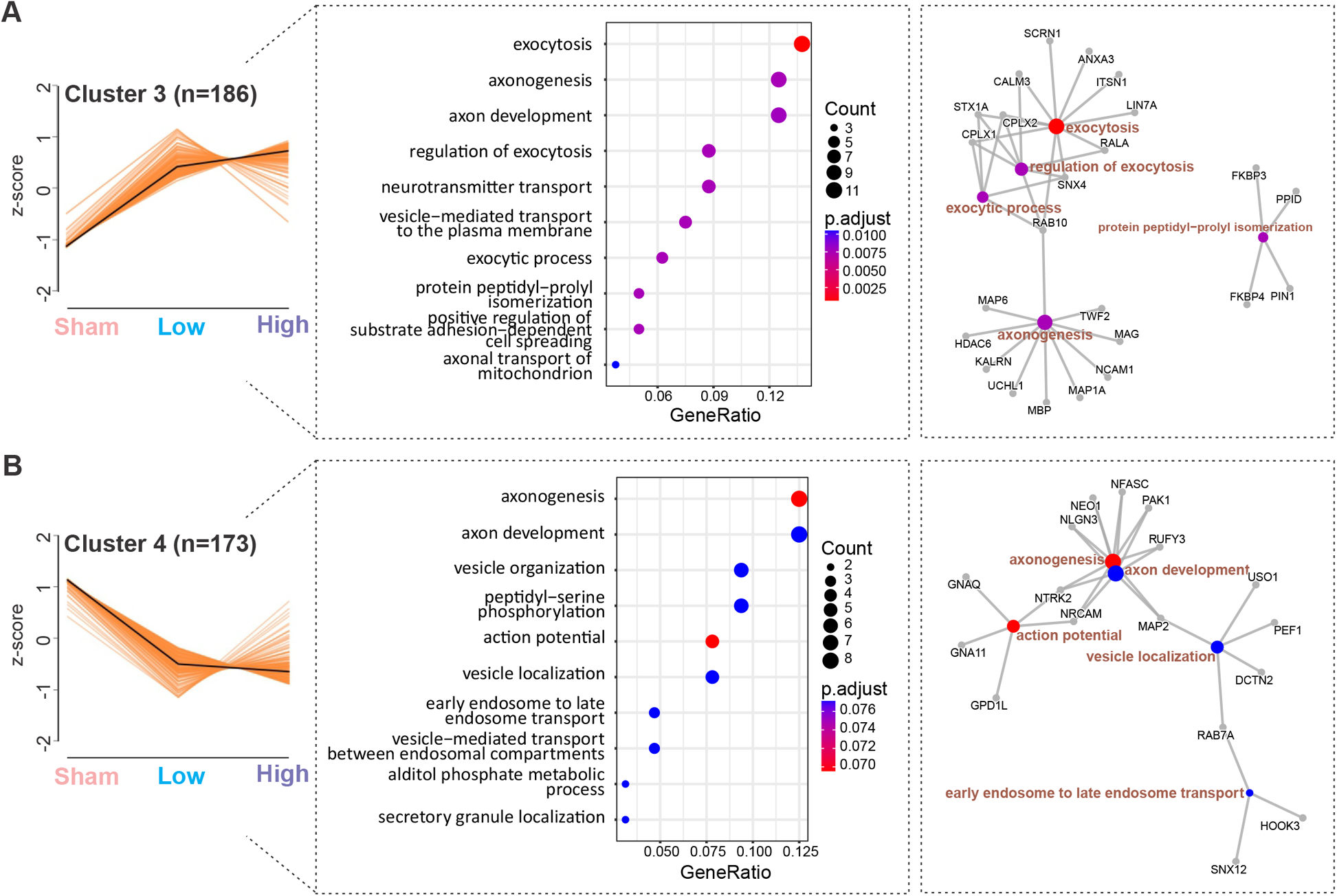
Treatment-frequency independent changes in the proteome. (A) Cluster 3 is defined by a similar pattern of increase in proteome triggered by ultrasound application. The cluster is represented by exocytosis and axon-associated processes. (B) Cluster 4 is characterized by a decrease in proteins following ultrasound application, but independent of the treatment regime. The processes associated with this cluster are related with axonal biology and vesicle localization.

**Supplementary Figure 2.**
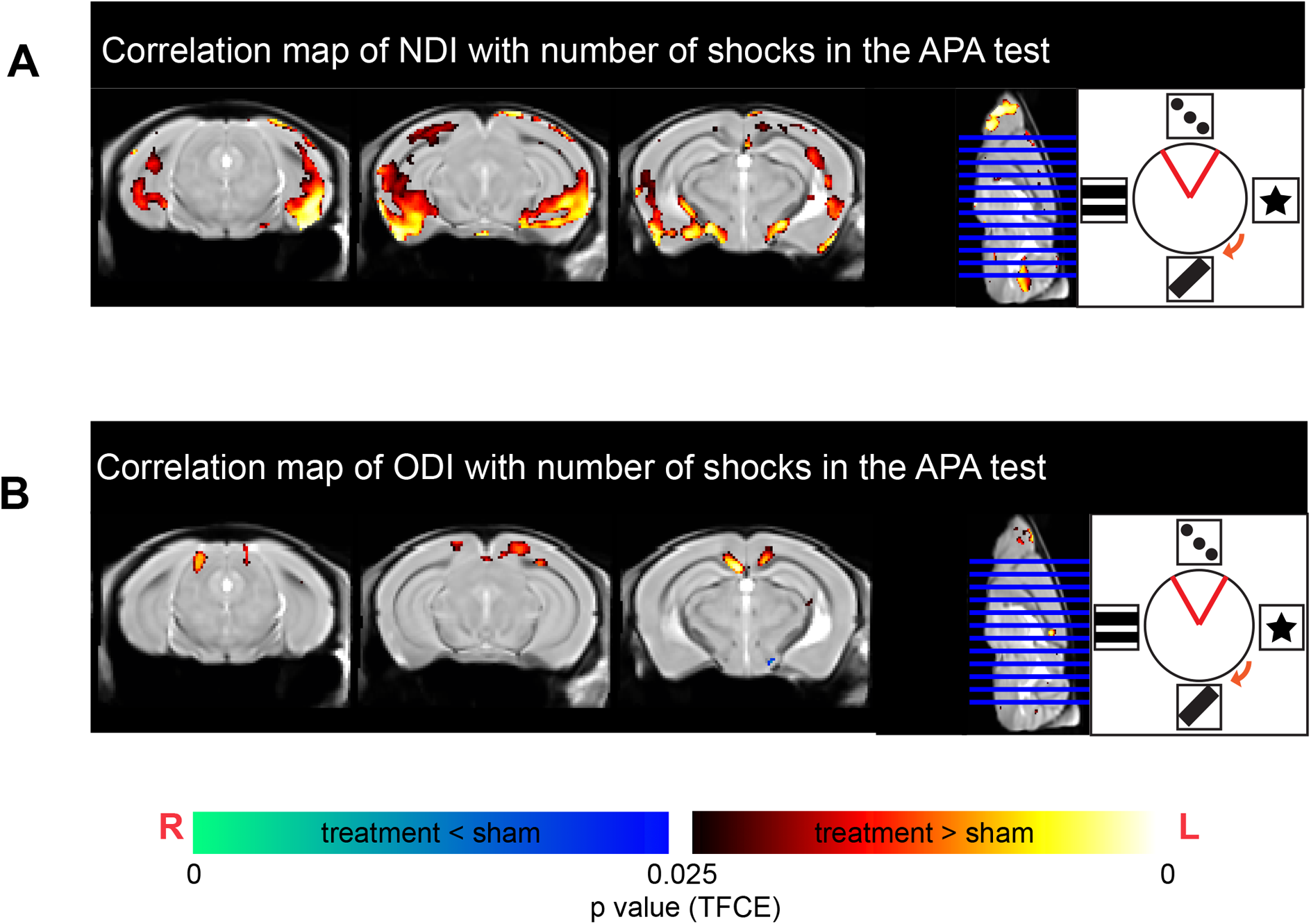
Diffusion MRI metrics correlated with performance in the APA test. (A) Neurite dispersion index (NDI) is positively correlated with number of shocks received in the APA test in ultrasound-treated mice. (B) Orientation dispersion index (ODI) was positively correlated with number of shocks received in the APA test in ultrasound-treated mice, with higher ODI in the corpus collosum being correlated with better performance in the APA (Heat map indicates voxels with p< 0.05 correlation. Indicative coronal and mid-sagittal slices displayed).

